# Sequence-Driven Drug-Target Affinity Prediction Via Graph Attention Networks and Bidirectional Cross-Attention Fusion

**DOI:** 10.64898/2026.04.03.716294

**Authors:** Zakeerhussain D Kudari, Virendra Singh Kaira, P Sai Shankar, Ruchika Bhat, G. Jaiganesh

## Abstract

Accurate prediction of drug-target affinity (DTA) is a core challenge in computational drug discovery. Structure-based methods depend on experimentally determined protein coordinates, which are unavailable for most drug-relevant targets. sequence-only approaches, in turn, operate on linear residue representations and lack an explicit mechanism to encode the spatial proximity relationships that govern protein-ligand interactions. We present XAttn-DTA, a sequence-driven framework that addresses both limitations without requiring experimental structural data. Drug molecules are encoded as 2D molecular graphs via multilayer Graph Attention Networks (GATs), capturing atomic topology and bond-level chemistry. Proteins are represented as residue-level graphs constructed from ESM2-predicted contact maps, that captures inter-residue coevolutionary and structural signals embedded within the sequence. The bidirectional cross-attention fusion module projects both embeddings into a shared latent space and applies dual multi-head cross-attention. This enables ligand and protein residue environments to inform one another. On the Davis benchmark, XAttn-DTA achieves a concordance index (CI) of 0.907 and MSE of 0.175, improving CI by 1.8% and reducing MSE by 9.3% over the strongest baseline. On KIBA, it achieves an MSE of 0.121, a 13.6% reduction. Under three strict cold-start settings across Davis, KIBA, and BindingDB, the model yields MSE reductions of up to 79.0% and CI improvements of up to 31.5% over the strongest baseline, demonstrating strong generalization to unseen scaffolds and novel protein families.

## 1. Introduction

Identifying protein-ligand interactions early on has long been one of the hardest problems in drug discovery, most candidates fail during clinical trials, even as screening technologies keep improving [1] [2] [3]. Structure-based methods like molecular docking and molecular dynamics simulations hit a familiar wall. They depend on experimentally resolved protein structures, struggle to handle large chemical libraries, and simply cannot help when high-quality structural data is unavailable, which remains the case for many therapeutically important proteins [1] [4] [5]. Together, these limitations push the field toward alternatives that work directly from sequence or independent of protein structures and learn from data rather than relying on resolved structures.

The increasing availability of biochemical interaction data, together with advances in machine learning, has led to the rapid adoption of data-driven approaches for modeling drug-target interactions [6]. Quantitatively predicting how strongly a drug binds to its target is typically expressed as a dissociation constant (*K*_*d*_), inhibition constant (*K*_*i*_), or half-maximal inhibitory concentration (*IC*_50_) has become one of the core regression tasks driving modern computational drug discovery [7].

Early computational approaches to drug-target affinity (DTA) prediction relied primarily on classical machine learning models, in which drugs and proteins were represented by handcrafted descriptors [8]. Many of these methods formulated DTA prediction as a binary classification task, aiming to determine whether a drug binds to a given target, rather than estimating binding strength [9]. Representative methods, such as this score, are computed by optimizing consistency across the original assay types using a statistical fusion procedure, KronRLS-MKL [10], NRLMF [11], and SELF-BLM [12], which employ kernel-based learning, neighborhood regularized matrix factorization, and ensemble learning with SVMs to predict interaction classes. While effective in certain settings, such formulations are limited in their ability to support high-precision drug design, where quantitative affinity estimation is essential.

DeepDTA [13] and WideDTA [14] were among the first to apply CNN [15] encoders directly to raw SMILES strings and protein sequences, bypassing manual feature engineering. While effective as baselines, encoding drugs as character sequences discards graph topology, and the learned representations are constrained by the expressive limits of linear sequence models. DeepConv-DTI [16], DeepAffinity [17], CO-VAE [18], and DeepLSTM [19] use recurrent architectures, variational autoencoders, and position-specific scoring matrices to enhance sequence-level representations of drugs and proteins.

Rather than encoding drugs as SMILES strings, GraphDTA [20] represented each molecule as a graph of atoms and bonds, processed by GNNs. CNNs handled protein sequences as before. The molecular graph representation alone was sufficient to produce consistent benchmark gains over SMILES-based encoders. Protein sequences continued to be handled by CNNs, nevertheless, the graph-based drug representation yielded consistent benchmark gains over SMILES-based encoders. PADME [21] and MGraphDTA [22] extended this direction, investigating graph architectures and dense connectivity patterns to better capture both local and global molecular structure. Constructing protein graphs at the atomic level requires significant computational power, whereas residue-level chains miss important details. Building useful protein graphs from sequence data alone, without experimental structures, remains a challenge.

Predicted contact maps offer a route to structurally informed protein representations without experimental coordinates. Each entry encodes the probability that two residues are spatially proximate in the folded state. Several contact predictors have been developed, such as RaptorX-Contact [23], DNCON2 [24], SPOT-Contact [25], DeepContact [26], DeepConPred [27], MetaPSICOV [28], and CCMpred [29]. However, many of these require significant computational resources when used at scale. DGraphDTA [30] showed that using a 0.5 threshold for ESM2 contact probabilities creates a sparse residue-level graph, which strikes a good balance between structural coverage and computational efficiency. Zheng et al. [31] approached drug-protein interaction prediction as a visual question-answering task and used two-dimensional distance maps to represent protein structure. GEFA [32] took a complementary approach, augmenting contact-derived protein graphs with language model node embeddings.

Transformer-based pretraining has substantially improved contextual representations for both molecular and protein inputs, notably through models such as Transformers [33], BERT [34], ESM [35], ChemBERTa [36], and ProteinBERT [37]. AttentionDTA [38] was one of the first models to use attention weighting for affinity prediction, which helped identify important interaction subsequences. Later models like TransformerCPI [39], CoaDTI [40], ML-DTI [41], MGPLI [42], and HyperAttentionDTI [43] built on this by adding mutual attention and multi-head mechanisms. However, most still encode each modality independently before fusion, leaving the problem of jointly conditioning each modality’s representation on the other largely unaddressed.

Despite significant progress, two primary challenges persist in sequence-based drug–target affinity (DTA) prediction. The first challenge is to build protein representations from sequence data that retain important interaction-related structures without using explicit three-dimensional coordinates. The second challenge is to combine different drug and protein representations so the model can handle new compounds and targets, especially in cold-start situations.

To resolve these challenges, we propose a sequence-driven DTA prediction framework that unifies three complementary directions, graph-based molecular modelling, sequence-derived protein structural representation, and attention-based interaction learning. Concretely, our contributions are as follows:

Rather than relying on experimentally resolved coordinates, we build residue-level protein graphs whose connectivity comes entirely from ESM2-predicted contact maps. Earlier contact-map-based approaches relied on relatively shallow predictors, which tend to miss the subtler, long-range relationships that actually influence protein folding and interactions. The contact probabilities predicted by ESM2 are influenced by both coevolutionary patterns and structural tendencies learned during pre-training on hundreds of millions of sequences. Node features integrate physicochemical residue properties with predicted secondary structure, enabling the generation of biologically meaningful representations even for proteins lacking experimentally resolved structures.

Instead of concatenating the two embeddings and passing the result to the regression head, both drug and protein representations are mapped into a common latent space. A bidirectional cross-attention module then operates across both modalities, allowing each representation to access the other’s global context before prediction.

The model is evaluated on three strict cold-start splits, where drugs, targets, and drug-target pairs are held out separately across Davis, KIBA, and BindingDB. The same splits are applied to four baseline models to ensure a fair comparison. The results show consistent MSE reductions and CI improvements over the best-performing baseline in the cold-start settings, which shows that sequence-derived structural representations with cross-modal fusion can provide stronger generalisation than the evaluated alternatives under cold-start conditions.

Evaluation on Davis and KIBA under warm-start and BindingDB, along with these datasets under three cold-start protocols, shows that the proposed framework achieves competitive performance under standard conditions and improved generalizations in the majority of held-out settings. These results support the viability of sequence-derived structural representations and cross-modal attention for sequence-based DTA prediction.

## 2. Materials and Methods

Our framework predicts binding affinity by jointly encoding drug molecular graphs and sequence-derived protein graphs, fusing their representations via bidirectional cross-attention, and regressing the resulting representation through a multilayer perceptron. The overall pipeline is illustrated in Figure 1.

**Figure 1.**
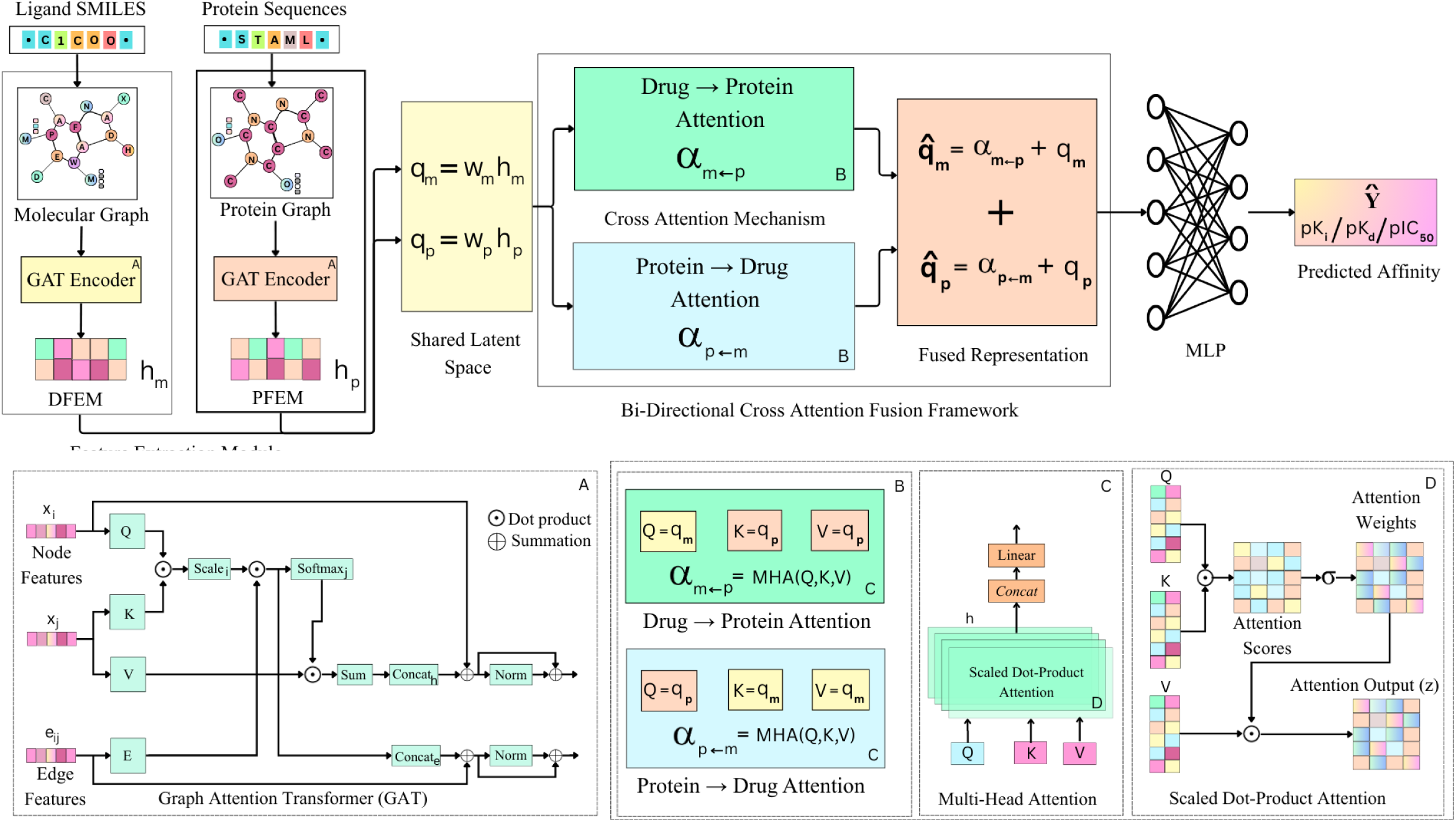
The framework is composed of four main components. (1) Drug Feature Extraction Module (DFEM): The SMILES string is converted into a molecular graph and encoded using stacked GAT layers, followed by graph-level pooling to obtain the drug embedding (ℎ_*m*_). (2) Protein Feature Extraction Module (PFEM): The amino acid sequence is converted into a protein graph and processed through GAT layers to produce the protein embedding embeddings (ℎ_*p*_).(3) Bidirectional Cross-Attention Fusion: The embeddings ℎ_*m*_ and ℎ_*p*_ are projected into a shared latent space and interact through dual multi-head cross-attention mechanisms, enabling the molecule to attend to the protein and the protein to attend to the molecule. This produces a fused representation capturing cross-modal dependencies. (4) Prediction Module: The fused vector is passed through a multilayer perceptron to regress the final binding affinity.

### 2.1 Dataset

We evaluate our model on two widely used drug-target affinity (DTA) benchmark datasets, the Davis dataset [44] and the KIBA dataset [45]. Both are standard benchmarks for assessing machine learning models in protein-ligand interaction prediction.

#### 2.1.1 Davis Dataset

The Davis kinase dataset [44] includes interaction measurements for 68 inhibitors and 442 kinase targets, with binding strength measured as *K*_*d*_. Before training, we apply a log transformation to the raw *K*_*d*_ values to compress the dynamic range and reduce skewness in the regression targets [13] [46]. As is standard [47], we convert these values to a log scale *pK*_*d*_using

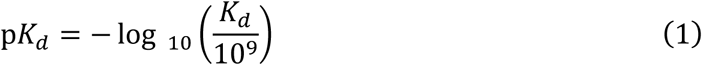

The SMILES of the small-molecule inhibitors have a median length of 55 characters (range: 32-81). Protein sequences were collected from UniProt [48], with a coevolutionary sequence length of 745 (range:244 to 2549).

#### 2.1.2 KIBA Dataset

The KIBA dataset [45] consolidates Ki, Kd, and IC50 measurements from multiple assay sources using a statistical model designed to maximize cross-assay consistency. This approach produces a unified, continuous bioactivity score for each drug-target pair. We adopt the processed version introduced by [47], which filters the dataset to include 2,068 ligands and 229 proteins, ensuring each appears in at least 10 measured interactions. Following [13] [47], KIBA scores are calculated as follows:

i. for each KIBA score, its negative was taken,
ii. the minimum value among the negatives was chosen and
iii. the absolute value of the minimum was added to all negative scores, thus constructing the final form of the KIBA scores.

SMILES strings have a median length of 58 characters (range: 20-590). Protein sequences were sourced via UniProt identifiers, with a median sequence length of 730 (range:215 to 4128). We remove all duplicated drug-target entries and ensure consistent SMILES and sequence mappings across both datasets. Invalid, missing, or corrupted molecular and sequence records are filtered out.

#### 2.1.3 BindingDB Dataset

BindingDB [49] It is a large, publicly available database of experimentally measured binding affinities encompassing interactions between small molecules and protein targets. Davis and KIBA are focused benchmarks. Davis uses curated kinase interactions, and KIBA uses composite bioactivity scores. In contrast, BindingDB collects binding measurements for a wider variety of target families and a more diverse set of chemicals. Its targets include enzymes, G protein-coupled receptors (GPCRs), ion channels, and nuclear receptors from different therapeutic areas. Before training, raw *K*_*d*_ values are log-transformed to *pK*_*d*_ as we did in the DAVIS dataset. BindingDB is included exclusively for cold-start evaluation.

The SMILES of the small-molecule inhibitors have a median length of 64 characters (range: 46-914). Protein sequences were collected from UniProt [48], with a median sequence length of 739 (range:86 to 2671). The final dataset statistics used for our experiments are summarized in Table 1.

**Table 1.** Summary statistics of the Davis and KIBA datasets used in our experiments.

| Datasets | Number of Compounds | Number of Proteins | Number of Interactions | Affinity Type |
| --- | --- | --- | --- | --- |
| Davis | 68 | 442 | 25772 | $K_d$, transformed to $pK_d$ |
| KIBA | 2068 | 229 | 118254 | KIBA score ($K_i$) |
| BindingDB | 42236 | 1088 | 9887 | $K_d$, transformed to $pK_d$ |

### 2.2 Graph Representation

Drug and protein inputs are both represented as graphs to preserve the relational structure that linear encodings or fixed-length descriptors discard. A drug molecule is represented as *Gm* = (*Vm*, *Em*), where *Vm* is the set of heavy atoms and *Em* the set of covalent bonds. A protein is represented as *G*_*p*_ = (*V*_*p*_, *E*_*p*_), where *V*_*p*_ contains one node per residue, and *E*_*p*_ encodes predicted residue-residue contacts. Neither graph requires crystallographic or cryo-EM input, as both are derived entirely from primary sequence data.

#### 2.2.1 Drug graph construction

Each atom node *i* in *V*_*m*_is assigned a feature vector *x*_*i*_ =∈ ℝ^(*F*^*a*+ *Fm*), which concatenates *L*1-normalised atom-level descriptors (as detailed in Table 2) with eight molecule-level physicochemical properties: LogP, HBA, HBD, aliphatic carbon count, aromatic carbon count, TPSA, volume, and aromatic ring count. These properties are broadcast uniformly to all nodes. Each bond (ⅈ, *j*) ∈ *E*_*m*_ is associated with a scalar weight that encodes the bond type (single, aromatic, double, or triple). Self-loops are incorporated to preserve node identity during message passing. The resulting graph is defined as *G*_*m*_ = (*X*, *Em*, *W*), where *X* ∈ ℝ^*N*^ ^×(*F*^*a*+ *Fm*), where *W* ∈ ℝ^|*E*^*m*|, and *N* denotes the number of atoms in the molecular graph, that is, *N* = |*V*_*m*_|.

**Table 2.** Classification of atom-level and molecule-level descriptors used in the drug graph encoder.

| Feature Name | Description | Dimensionality |
| --- | --- | --- |
| Atom Type | One-hot encoding of an atomic symbol (C, N, O, S, ...) | 44 |
| Atom Neighbors | One-hot encoding of atom degree (0–10) | 11 |
| Number of Hydrogens | Number of attached hydrogens | 11 |
| Atom is_Aromatic | Aromaticity flag (binary) | 01 |
| Atomic Weight | Normalized atomic mass. | 01 |
| Positive Charged | Atom has a positive formal charge | 01 |
| Negative Charged | Atom has a negative formal charge | 01 |
| Hybridization | Encodes sp / sp<sup>2</sup> / sp<sup>3</sup> hybridization | 01 |
| Formal Charge | Integer atom charge | 01 |
| LogP | Lipophilicity descriptor | 01 |
| HBA | Hydrogen Bond Acceptors Molecule-level count | 01 |
| HBD | Hydrogen Bond Donors Molecule-level count | 01 |
| Aliphatic Carbons | Count of aliphatic carbon atoms | 01 |
| Aromatic Carbons | Count of aromatic carbon atoms | 01 |
| TPSA | Topological Polar Surface Area | 01 |
| Volume | Approximate molecular volume | 01 |
| Number of Aromatic Rings | Aromatic subset of rings | 01 |

Figure 2 shows that the workflow for constructing drug molecular graphs from SMILES strings. A SMILES string is first processed to identify atomic and bonding information, which is then used to generate the molecular graph. Node and edge features are subsequently computed to produce the final graph representation used for learning.

**Figure 2.**
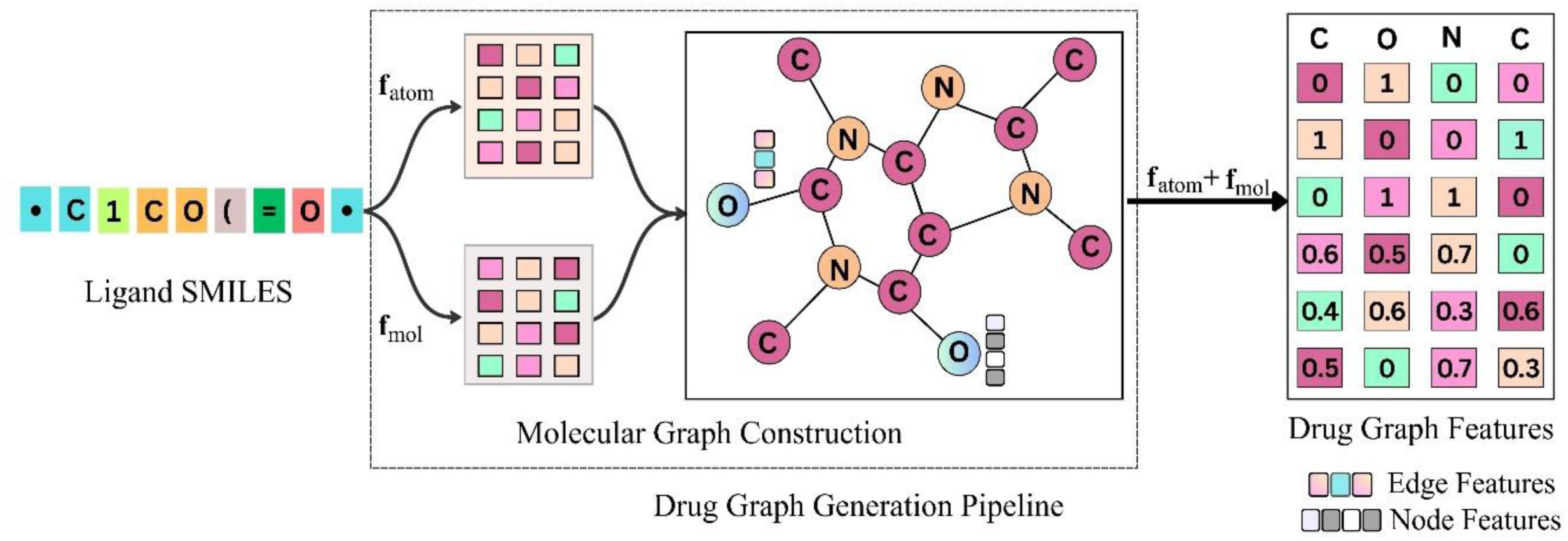
The workflow for constructing drug molecular graphs from SMILES strings. A SMILES string is first processed to identify atomic and bonding information, which is then used to generate the molecular graph. Node and edge features are subsequently computed to produce the final graph representation used for learning.

#### 2.2.2 Protein Graph Construction

Given a protein sequence of amino acids, *s* = (*a*_1_, *a*_2_, …, *a*_*N*_*p*), the residue-residue contact probability matrix *C*[0,1]^*N*^*p*×*Np* is generated using ESM2 [50]. Self-contacts are assigned *C*_*ii*_ = 1, and positions beyond the actual sequence length are masked. If the predicted contact score *C*_*ij*_ is greater than 0.50 [23], the edge is generated between residues *i* and *j*, which represents the structural informativeness. The edge weight *w*_*ij*_ = *C*_*ij*_ ∈ [0,1] represents the model’s confidence in spatial proximity, resulting in the weighted edge set

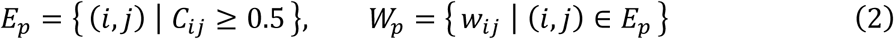

Encoding each residue node *i* with a feature vector *x*_*i*_ ∈ ℝ^*F*^*p* with feature parameters such as residue identity, hydrophobicity scales, solvent exposure, and predicted secondary structure, as summarized in Table 3. This results in the graph representation G𝒢_*p*_ = (*X*_*p*_, *E*_*p*_, *W*_*p*_), where *X*_*p*_ ∈ ℝ^*N*^*p*×*Fp*. As shown in Figure 3 which depicts the two-branch pipeline, where ESM2 generates the contact map concurrently with a processing module that computes residue-level features.

**Figure 3.**
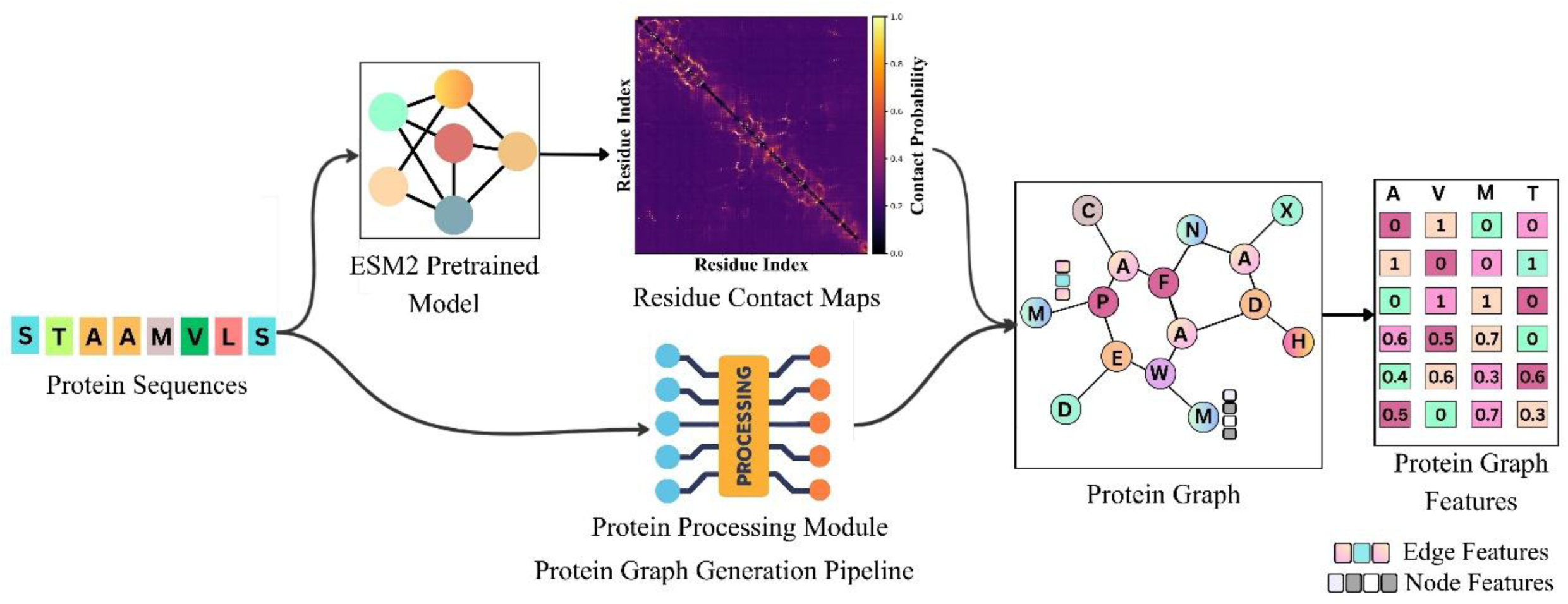
Two branch architecture for protein graph generation. The input protein sequence is first processed by ESM2 to predict residue contact maps, while a parallel protein processing module produces residue specific feature vectors. The contact information and residue features are integrated to construct the final protein graph

**Table 3.** Residue-Level and physicochemical features used for protein graph construction.

| Feature Name | Description | Dimensionality |
| --- | --- | --- |
| Residue Type | One-hot encoding of amino acid identity (20 std + unknown) | 21 |
| Aliphatic Residues | Indicator if residue is aliphatic (A, I, L, M, V) | 1 |
| Aromatic Residues | Indicator if residue is aromatic (F, W, Y) | 1 |
| Acidic Residues | Indicator for acidic amino acids (D, E) | 1 |
| Basic Residues | Indicator for basic amino acids (K, R, H) | 1 |
| Neutral Residues | Indicator for neutral residues | 1 |
| Residue Weights | Normalized residue molecular weight | 1 |
| pKa (-COOH group) | Dissociation constant of the acidic side | 1 |
| pKb (-NH<sub>3</sub> group) | Dissociation constant of the basic side | 1 |
| pKx (another group) | Dissociation constant of additional ionizable groups | 1 |
| Isoelectric Point (pI) | pH at which the residue carries no net charge | 1 |
| Hydrophobicity at pH = 2 | Low-pH hydrophobicity scale | 1 |
| Hydrophobicity at pH = 7 | Neutral-pH hydrophobicity scale | 1 |
| Kyte-Doolittle Hydropathy | Standard hydropathy index | 1 |
| Hopp-Woods Hydropathy | Alternative hydropathy scale for antigenicity | 1 |
| Relative Solvent Accessibility (RSA) | Normalized per-residue solvent exposure | 1 |
| Secondary Structure (helix, sheet, coil) | Trinary structure encoding from predictors | 3 |
| Aliphatic Index | Contribution of aliphatic side chains (per residue or normalized) | 1 |

Two-branch architecture for protein graph generation is shown in Figure 3. ESM2 processes the input protein sequence to predict residue contact maps, while a parallel protein processing module generates residue-specific feature vectors. The contact information and residue features are then integrated to construct the final protein graph.

### 2.3 Graph Attention Encoders

Both *G*_*m*_ and *G*_*p*_ are encoded by the same multi-layer GAT architecture, instantiated with independent parameters for each modality. Let 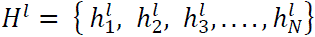, denote node features at layer l, with *H*^(0)^ = *X*_*m*_ (drug) or *H*^(0)^ = *X*_*p*_ (protein).

For each edge (*i*, *j*), a per-head attention score is computed by a linear scoring function with LeakyReLU nonlinearity (negative slope 0.2) applied to the pairwise concatenation of projected node representations. Raw scores are converted to a probability distribution over each node’s neighbourhood via softmax normalisation:

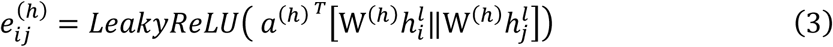

where a^(ℎ)^ ∈ ℝ^2*F*^′ is a learnable parameter vector and ‖ denotes concatenation.

Scores are computed exclusively over directly connected neighbours *j* ∈ 𝒩_*i*_, and normalised within each neighbourhood via softmax, yielding attention coefficients that sum to one over each local neighbourhood.

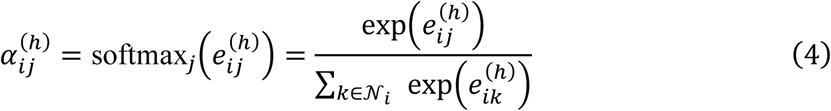

Each attention head h independently aggregates neighboring features based on its learned coefficients.

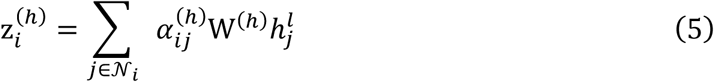

The outputs of H independent heads are concatenated to form the updated node representation z ∈ ℝ^*HF*^′, thereby preserving complementary structural information learned in parallel across the heads.

The concatenated representation is passed through layer normalization, followed by *ReLU* activation. A residual connection is applied by projecting the input features 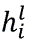 to match the output dimensionality via W_skip_ ∈ ℝ^(*HF*^ ^×^ ^F)^ and adding the result

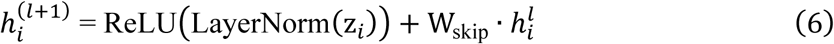

Skip connections preserve input information and stabilize training in deeper architectures. After L layers, a graph-level embedding is obtained via mean pooling over all atom nodes

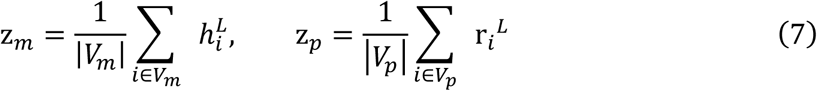

A two-layer MLP with a residual connection refines this representation

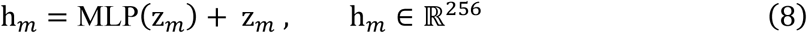

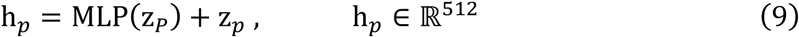

The drug embedding (h_*m*_) represents atomic neighbourhoods, chemically meaningful substructures, and global molecular topology. This embedding is subsequently provided to the bidirectional cross-attention fusion module for interaction-level modelling. The protein embedding (h_*p*_) captures residue-level biochemical properties, contact-derived structural topology, and both short- and long-range inter-residue dependencies.

### 2.4 Fusion Framework

After independently encoding the drug and protein graphs, we introduce a cross-modal fusion module that models interaction-level dependencies between the two modalities. Instead of directly fusing representations, this module projects both embeddings into a shared latent space and applies bidirectional multi-head attention, allowing each modality’s representation to be selectively updated by the other’s context before final concatenation and regression.

#### 2.4.1 Shared Latent Space

The embeddings of the drug and protein have different dimensionality, h_*m*_ ∈ ℝ^256^ and h_*p*_ ∈ ℝ^512^, respectively. In order to compute attention across modalities, embeddings are projected into a shared latent space of dimension d via independent linear transformations, yielding q_*m*_, q_*p*_ ∈ ℝ^*d*^.

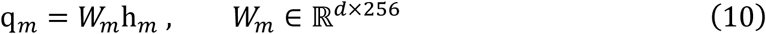

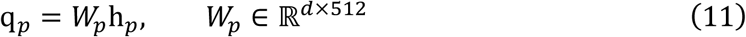

Both vectors are reshaped to 1 × *d* token sequences to serve as valid inputs to multi-head attention layers. This projection aligns the two modalities dimensionally while allowing each to retain its own representational structure prior to interaction modelling.

#### 2.4.2 Bidirectional Cross-Attention

To integrate the mutual information between the drug and protein at the global level, we apply bidirectional multi-head attention over their projected representations, alternating the roles of the modalities as query and key-value sources. The drug embedding attends to the protein’s aggregate biochemical context, and the protein embedding attends to the drug’s aggregate chemical profile.

Molecule attends to protein: The drug projection q_*m*_ acts as a query, while q_*p*_ provides keys and values, producing a protein-conditioned update to the drug representation.

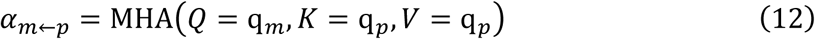

Protein attends to molecule: The protein projection q_*p*_ acts as a query, while q_*m*_ provides keys and values, producing a drug-conditioned update to the protein representation.

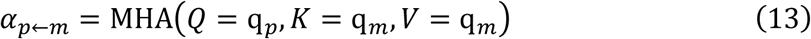

Each cross-attention output is stabilised via a residual connection followed by layer normalisation, following standard Transformer practice [33]

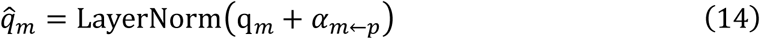

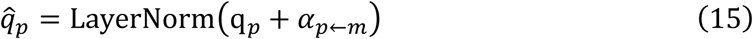

This attention mechanism operates over globally pooled drug and protein embeddings rather than over individual atom or residue tokens.

Consequently, cross-modal conditioning is performed at the level of aggregated representations, allowing the model to assess whether and to what extent the overall chemical characteristics of a drug complement the protein’s global biochemical environment. However, this formulation fails to capture fine-grained interaction details, such as specific atom-residue contacts that frequently determine binding specificity. This design trade-off reduces model complexity and circumvents the quadratic computational cost of token-level cross-attention, but at the expense of interaction-site resolution.

#### 2.4.3 Joint Fusion Representation

The cross-attended embeddings are squeezed to d-dimensional vectors and combined with their original projections via residual addition before concatenation, reinforcing interaction-specific signals with the unmodified modality context

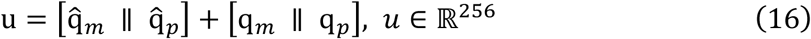

where ∥ denotes vector concatenation. The resulting fused embedding *u* ∈ ℝ^256^ encodes bidirectional cross-modal dependencies between drug substructures and protein residue environments, and serves as the sole input to the prediction module.

### 2.5 Prediction Module

The fused representation *u* ∈ ℝ^256^ is passed through a three-layer MLP to regress the final binding affinity score. The prediction head applies ReLU nonlinearities and dropout regularisation (rate *p*=0.2, applied during training only) after each hidden layer. DTA prediction is formulated as a regression task. The model is trained by minimising the mean squared error (MSE) between predicted and ground-truth affinity values over a training set of n drug-protein pairs:

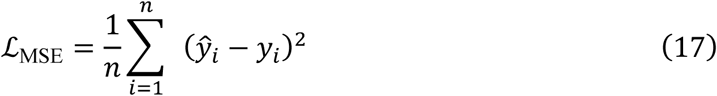

where *y*^_*i*_is the predicted affinity and *y*_*i*_is the experimentally measured ground truth (expressed as in *K*_*i*_, *K*_*d*_ or *IC*_50_in negative log scale). MSE is chosen for its convexity and its well-established use in continuous affinity regression benchmarks [30][41].

### 2.6 Evaluation Metrics

We evaluate model performance using three metrics and standards in drug-target affinity prediction [43], concordance index (CI), root mean squared error (RMSE), and mean squared error (MSE). CI quantifies ranking performance while MSE and RMSE quantify regression accuracy.

CI [51] is a pairwise ranking metric that evaluates the model’s ability to correctly order interaction strengths. For any two drug–target pairs with differing true affinities, CI measures the proportion of cases in which the model assigns a higher predicted score to the more potent binder. A CI value of 1.0 indicates perfect ranking, whereas a value of 0.5 corresponds to performance expected under random prediction.

Formally:

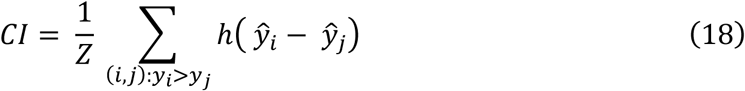

where *y*^_*i*_ and *y*^_*j*_ are predicted affinities for a pair with true affinities *y*_*i*_ > *y*_*j*_, h(·) is the Heaviside step function ℎ(*x*) = {1 *if x* > 0; 0.5 *if x* = 0; 0 *if x* < 0}, and *Z* is a normalisation constant equal to the total number of qualifying pairs.

MSE measures the mean squared deviation between predicted and experimentally measured affinity values

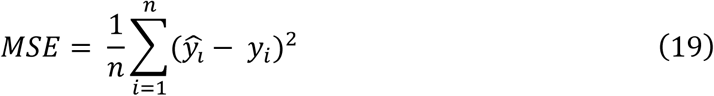

where *y*_*i*_ is the ground-truth affinity for the *i*-th drug-protein pair, *y*^_*i*_ is the model prediction, and *n* is the number of samples. Lower MSE indicates higher numerical accuracy.

RMSE is the square root of MSE, returning the error to the same units as the affinity target and facilitating direct comparison with the scale of measured values

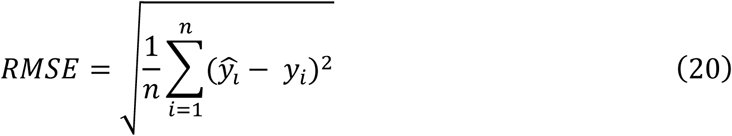

Together, CI and MSE/RMSE provide complementary assessments, such as CI captures the model’s ability to correctly prioritise compounds by relative binding strength, while MSE and RMSE quantify absolute prediction accuracy.

## 3. Results and Discussion

### 3.1 Experimental Setup

All experiments used in this process are from the Davis, KIBA, and BindingDB datasets described in the dataset section. We clean the dataset by transforming the target value as in the case of Davis and BindingDB, samples with duplicate entries, invalid SMILES strings, or incomplete protein sequences are removed prior to training. Drug molecules are converted into molecular graphs using RDKit, and protein contact maps are predicted from amino acid sequences using ESM2.

For warm-start evaluation, we follow standard practice and apply an 80/10/10 train/validation/test split. For cold-start evaluation, three additional splits are constructed:

i. drug cold start, where no test compound appears in training
ii. target cold-start, where no test protein appears in training, and
iii. drug–target cold-start, where neither drugs nor proteins in the test set are seen during training. Cold-start splits allocate 70%, 10%, and 20% of unique entities to training, validation, and test partitions respectively, following the protocol [9].

The model is trained with an initial learning rate of 5×10⁻³, Adam as the optimizer, and a batch size of 64. Dropout is applied to all MLP layers at a rate of p = 0.2. Drug embeddings have dimensionality 256 and protein embeddings 512, while both are projected to a shared 128-dimensional latent space for cross-attention. Each GAT [52] encoder uses 4 attention heads per layer. Early stopping with a patience of 20 epochs is applied based on validation MSE. All experiments and code are implemented using PyTorch and executed on an NVIDIA RTX 6000 Ada (48 GB). Hyperparameter search ranges and selected values are reported in Table 4.

**Table 4.** Hyperparameter and search ranges used in model tuning.

| Hyperparameter | Setting | Defaults |
| --- | --- | --- |
| Learning rate | (1e-3, 5e-3, 1e-4, 5e-4) | 5e-3 |
| Number of GAT Layers | (2, 3, 4, 5) | 3 |
| Number of Attention Heads | (2, 4, 8) | 4 |
| Number of Layers in MLP | (2, 3, 4, 5) | 3 |
| Dropout | (0.20, 0.30) | 0.20 |
| Batch Size | (32, 64, 128, 256, 512) | 64 |
| Epochs | (100, 200, 300, 1000) | 300 |
| Patience | (10, 20, 30) | 20 |

### 3.2 Performance Evaluation

Two separate baseline sets are used for warm-start, and three datasets are used for cold-start evaluation, reflecting differences in data availability across protocols and datasets. Performance is measured using mean squared error (MSE) and concordance index (CI).

For warm-start evaluation on Davis and KIBA, we compare against seven established methods DeepDTA [13], AttentionDTA [38], GraphDTA [20], TransformerCPI [39], ML-DTI [41], MGPLI [42] and AttentionMGT-DTA [9], for which published results under identical warm-start splits and metrics are available on both datasets.

For cold-start evaluation across Davis, KIBA, and BindingDB, a distinct set of four methods is used GraphDTA [20], DrugMGR [53], G-K BertDTA [54] and DTA-GTOmega [55]. This set differs from the warm-start baselines for a specific reason: while cold-start results for the warm-start baseline set could be obtained for Davis, published cold-start results for KIBA and BindingDB were unavailable for those models. Using only Davis cold-start results would produce an inconsistent baseline set across the three cold-start datasets, making cross-dataset comparisons unreliable. The four cold-start baselines were therefore selected as the models for which complete results across all cold-start settings and all three datasets are available, following the evaluation protocol of DTA-GTOmega [55].

These baselines span four architectural paradigms. Sequence-based encoders such as DeepDTA [12] use parallel CNNs over SMILES and protein sequences. AttentionDTA [38] adds attention over CNN-extracted features to identify interaction-relevant subsequences. Graph-based drug encoding models, GraphDTA [20], represent drugs as molecular graphs encoded by GNNs while processing proteins with CNNs. DrugMGR [53] extends this with dense connectivity and multigranular protein feature extraction, respectively. Transformer and multi-granular encoding model TransformerCPI [39] apply self-attention with label-reversal validation. ML-DTI [41] employs mutual multi-head attention, enabling each encoder to attend to the others. MGPLI [42] tokenizes both modalities at the character and segment level using SentencePiece before transformer encoding. Hybrid structure-aware models such as AttentionMGT-DTA [9] build protein pocket graphs from AlphaFold2 structures and fuse them with sequence embeddings via cross-attention GK-BertDTA [54] combines DenseNet protein features with graph iso orphism network drug encoding and KB-BERT semantic embeddings. DTA-GTOmega [55] constructs protein graphs from OmegaFold-predicted 3D structures and models atom-residue interactions via co-attention.

#### 3.2.1 Performance on benchmark datasets

The proposed model outperforms all seven baseline methods on both datasets, with the highest concordance index (CI) and lowest mean squared error (MSE). On the Davis dataset, it achieves a CI of 0.907 and an MSE of 0.175. On the KIBA dataset, it records a CI of 0.894 and an MSE of 0.121. Results on KIBA and Davis as reported in Table 5 and Figure 4. Baseline results are taken from AttentionMGT-DTA [9] and evaluated under identical data splits and metrics. On the two divergent datasets, the kinase-centric Davis dataset with a narrow chemical space, and KIBA which integrates heterogeneous bioactivity measurements spanning substantially broader target and ligand diversity, our model’s consistent performance suggests that the improvements are not dataset-specific artifacts but reflect a genuine advantage in representational learning. Our model improves CI by 1.8% and reduces MSE by 9.3%. For the Davis dataset relative to the strongest baseline (AttentionMGT-DTA: CI = 0.891, MSE = 0.193), against sequence-only baselines, MSE reductions are more pronounced 8.4% over DeepDTA, 10 3% over AttentionDTA, 10.7% over ML-DTI, and 19.7% over MGPLI.

**Figure 4.**
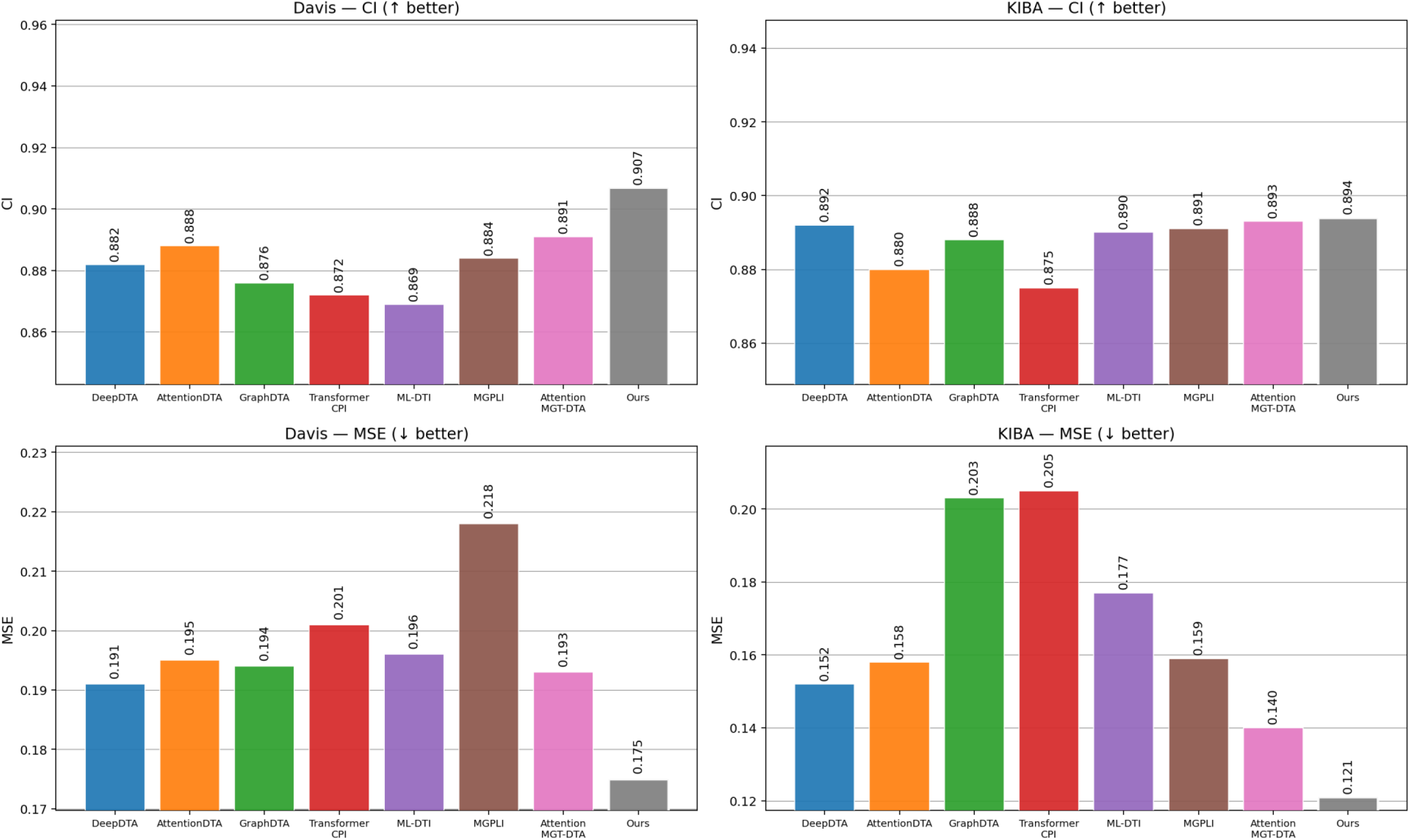
Comparative performance of the proposed model and baseline methods across benchmark datasets measured using MSE and CI. Lower MSE and higher CI indicate better predictive performance.

**Table 5.** Performance comparison between our model and baseline methods on the DAVIS and KIBA dataset.#.

| Dataset | Model | CI | MSE |
| --- | --- | --- | --- |
| Davis | DeepDTA<sup>a</sup> | 0.882 | 0.191 |
|  | AttentionDTA<sup>a</sup> | 0.888 | 0.195 |
|  | GraphDTA<sup>a</sup> | 0.876 | 0.194 |
|  | TransformerCPI<sup>a</sup> | 0.872 | 0.201 |
|  | ML-DTI<sup>a</sup> | 0.869 | 0.196 |
|  | MGPLI<sup>a</sup> | 0.884 | 0.218 |
|  | AttentionMGT-DTA<sup>a</sup> | 0.891 | 0.193 |
|  | Present Work | **0.907** | **0.175** |
| KIBA | DeepDTA<sup>a</sup> | 0.892 | 0.152 |
|  | AttentionDTA<sup>a</sup> | 0.880 | 0.158 |
|  | GraphDTA<sup>a</sup> | 0.888 | 0.203 |
|  | TransformerCPI<sup>a</sup> | 0.875 | 0.205 |
|  | ML-DTI<sup>a</sup> | 0.890 | 0.177 |
|  | MGPLI<sup>a</sup> | 0.891 | 0.159 |
|  | AttentionMGT-DTA<sup>a</sup> | 0.893 | 0.140 |
|  | Present Work | **0.894** | **0.121** |

The observed improvements over sequence-based encoders support the hypothesis that graph-structured drug representations capture atomic topology not fully recoverable by SMILES-based models, and that ESM2-derived contact maps provide interaction-relevant protein topology absent in linear sequence encoders. Notably, the CI range across all seven baselines on Davis is narrow (0.869-0.891), reflecting a general saturation of ranking performance at this benchmark. The more informative differentiation is therefore in MSE, where the model’s absolute reduction of 0.016 over the strongest baseline represents a meaningful improvement in regression accuracy at a performance level where marginal gains are increasingly difficult to achieve.

For Davis dataset relative to the strongest baseline (AttentionMGT-DTA: CI = 0.891, MSE = 0.193), the model improves CI by 1.8% and reduces MSE by 9.3%. Against sequence-only baselines, MSE reductions are more pronounced 8.4% over DeepDTA, 10.3% over AttentionDTA, 10.7% over ML-DTI, and 19.7% over MGPLI. These gains over sequence-based encoders are consistent with the hypothesis that graph-structured drug representations capture atomic topology that SMILES-based models cannot fully recover, and that ESM2-derived contact maps provide interaction-relevant protein topology absent from linear sequence encoders. Notably, the CI range across all seven baselines on Davis is narrow (0.869-0.891), reflecting a general saturation of ranking performance at this benchmark. The more informative differentiation is therefore in MSE, where the model’s absolute reduction of 0.016 over the strongest baseline represents a meaningful improvement in regression accuracy at a performance level where marginal gains are increasingly difficult to achieve.

The performance pattern on the KIBA dataset is qualitatively different and analytically important. The CI gain over the strong st baseline is marginal (0.894 vs 0.893, a 0.11% improvement). In contrast, the MSE reduction is substantially larger, 13.6% over AttentionMGT-DTA, 20.4% over DeepDTA, 23.4% over AttentionDTA, 31.6% over ML-DTI, and 41.0% over TransformerCPI. This asymmetry between CI and MSE gains on KIBA reflects a structural property of the benchmark rather than a model limitation. The CI range across all seven baselines on KIBA is only 0.018 (0.875–0.893), an even narrower spread than on Davis, indicating that affinity ranking is a near-solved task at this performance tier and that CI provides limited discriminative power among competitive models. In contrast, the MSE range across baselines on KIBA spans 0.065 (0.140–0.205), preserving meaningful separation. The model’s MSE of 0.121 falls below the entire baseline range, indicating that the improvement in absolute prediction accuracy is both statistically substantive and practically meaningful for applications requiring precise affinity estimation rather than relative ranking alone.

AttentionMGT-DTA [9] is the most architecturally relevant point of comparison, as it employs cross-attention fusion and constructs protein graphs from AlphaFold2-predicted pocket structures, giving it access to three-dimensional protein geometry that our model does not. Despite this structural advantage, XAttn-DTA outperforms AttentionMGT-DTA on MSE by 9.3% on Davis and 1.6% on KIBA, and matches or exceeds it on CI on both datasets. The comparison is especially telling, as both models share a graph-based drug encoder and cross-attention fusion, differing only in how the protein is represented. Where AttentionMGT-DTA builds pocket graphs from AlphaFold2 predicted structures, the proposed model derives contact maps directly from ESM2, with no structural prediction, no pocket annotation, and no three-dimensional coordinates. That substitution alone is sufficient to improve regression accuracy on both benchmarks. This finding supports the view that coevolutionary sequence signals captured by protein language models encode binding-relevant structural information that is competitive with, and in some respects complementary to, explicitly predicted three-dimensional geometry.

Examining the full baseline ranking reveals that a consistent trend model incorporating more expressive molecular representations tends to achieve lower MSE, even when CI gains are modest. Sequence-based models (DeepDTA, AttentionDTA, TransformerCPI) cluster at higher MSE values on both datasets, while graph-based drug encoders (GraphDTA, AttentionMGT-DTA) and the proposed model achieve progressively lower MSE. The improvement from GraphDTA to the proposed model on KIBA (MSE 0.203 to 0.121, a 40.4% reduction) quantifies the combined contribution of ESM2-derived protein graphs and bidirectional cross-attention fusion over standard GNN drug encoding with CNN protein encoding. While this comparison conflates multiple architectural differences, it establishes the magnitude of the overall representational advance relative to the graph-based DTA baseline.

While these results confirm competitive performance under standard evaluation conditions, warm-start benchmarks allow overlap between the training and test sets. Both drugs and proteins in the test set may have been observed during training in different interaction contexts. This can overstate real-world performance relative to deployment scenarios involving novel chemical scaffolds or protein families. The following section, therefore, evaluates the model under three strict cold-start settings that eliminate this overlap, providing a more reliable assessment of its generalisation capability.

#### 3.2.2 Performance on Cold-Start

Three strict cold-start protocols are used to probe generalisation, test drugs, test proteins, or both are withheld entirely from training. Table 6 and Figure 5 report results across Davis, KIBA, and BindingDB. For cold-start evaluation, we directly adopt the pre-defined splits provided by DTA-GTOmega [52], the same split protocol applied to Davis, KIBA and BindingDB in this work, ensuring that cold-start partitioning is consistent and directly comparable across all three benchmarks. As expected, all methods suffer performance degradation relative to warm-start conditions. Distributional shift is an inherent consequence of excluding entire scaffolds or protein families from the training set. The proposed model nonetheless records the lowest MSE and RMSE and the highest CI across the majority of these settings on both datasets.

**Figure 5.**
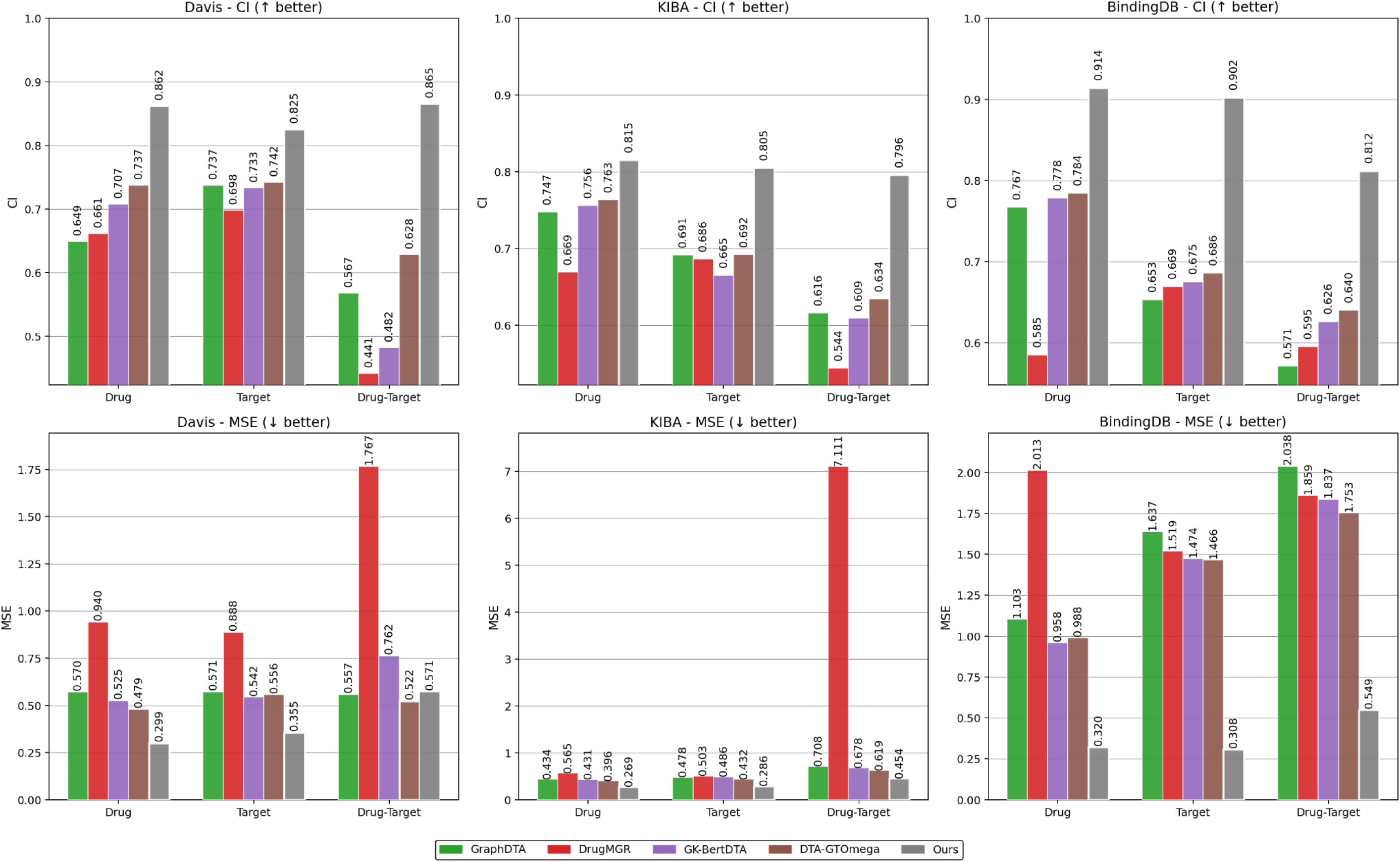
Comparative performance of the proposed model and baseline methods under cold-start settings (drug cold-start, target cold-start, and pair cold-start), evaluated using MSE and CI. Lower MSE and higher CI indicate better predictive accuracy and generalization ability.

**Table 6.** Performance comparison between our model and baseline methods on KIBA, Davis and BindingDB dataset under cold-start settings#.

| Settings | Models | KIBA |  |  | Davis |  |  | BindingDB |  |  |
| --- | --- | --- | --- | --- | --- | --- | --- | --- | --- | --- |
|  |  | MSE | RMSE | CI | MSE | RMSE | CI | MSE | RMSE | CI |
| Drug<br>Cold-start | GraphDTA<sup>*</sup> | 0.434 | 0.659 | 0.747 | 0.570 | 0.755 | 0.649 | 1.103 | 1.050 | 0.767 |
|  | DrugMGR<sup>*</sup> | 0.565 | 0.752 | 0.669 | 0.940 | 0.970 | 0.661 | 2.013 | 1.419 | 0.585 |
|  | G-K BertDTA<sup>*</sup> | 0.431 | 0.656 | 0.756 | 0.525 | 0.724 | 0.707 | 0.958 | 0.979 | 0.778 |
|  | DTA-GTOmega<sup>*</sup> | 0.396 | 0.629 | 0.763 | 0.479 | 0.692 | 0.737 | 0.988 | 0.994 | 0.784 |
|  | Present Work | **0.269** | **0.519** | **0.815** | **0.299** | **0.547** | **0.862** | **0.320** | **0.566** | **0.914** |
| Target<br>Cold-start | GraphDTA<sup>*</sup> | 0.478 | 0.692 | 0.691 | 0.571 | 0.756 | 0.737 | 1.637 | 1.279 | 0.653 |
|  | DrugMGR<sup>*</sup> | 0.503 | 0.709 | 0.686 | 0.888 | 0.942 | 0.698 | 1.519 | 1.233 | 0.669 |
|  | G-K BertDTA<sup>*</sup> | 0.486 | 0.697 | 0.665 | 0.542 | 0.736 | 0.733 | 1.474 | 1.214 | 0.675 |
|  | DTA-GTOmega<sup>*</sup> | 0.432 | 0.657 | 0.692 | 0.556 | 0.746 | 0.742 | 1.466 | 1.211 | 0.686 |
|  | Present Work | **0.286** | **0.535** | **0.805** | **0.355** | **0.596** | **0.825** | **0.308** | **0.544** | **0.902** |
| Drug-Target<br>Cold-start | GraphDTA<sup>*</sup> | 0.708 | 0.842 | 0.616 | 0.557 | 0.746 | 0.567 | 2.038 | 1.428 | 0.571 |
|  | DrugMGR<sup>*</sup> | 7.111 | 2.667 | 0.544 | 1.767 | 1.329 | 0.441 | 1.859 | 1.363 | 0.595 |
|  | G-K BertDTA<sup>*</sup> | 0.678 | 0.823 | 0.609 | 0.762 | 0.873 | 0.482 | 1.837 | 1.355 | 0.626 |
|  | DTA-GTOmega<sup>*</sup> | 0.619 | 0.787 | 0.634 | **0.522** | **0.722** | 0.628 | 1.753 | 1.324 | 0.640 |
|  | Present Work | **0.454** | **0.672** | **0.796** | 0.571 | 0.755 | **0.865** | **0.549** | **0.741** | **0.812** |

In the Drug Cold-Start Setting on Davis, our model achieves MSE = 0.299 and CI = 0.862, compared to the strongest baseline (DTA-GTOmega: MSE = 0.479, CI = 0.737), corresponding to a 37.6% reduction in MSE and a 17.0% improvement in CI. On KIBA, MSE = 0.269 and CI = 0.815, outperforming the best baseline (DTA-GT mega: MSE = 0.396, CI = 0.763) by 32.1% in MSE and 6.8% in CI. On BindingDB, yields the largest margins, CI = 0.914, reducing MSE by 66.6% relative to the strongest baseline (GK-BertDTA: MSE = 0. 58) and improving CI by 16.6% relative to DTA-GTOmega (0.784). The consistent advantages suggest that the GAT-based drug encoder is effectively learning a substructure representation that can generalize across various chemical compounds. It captures local atomic neighborhood patterns that are transferable, even when moving between different scaffold types. This means it’s not just memorizing the specific molecular graphs it was trained on, but rather developing a deeper understanding that applies more broadly in the chemical landscape.

Target cold-start results follow a similar pattern. On Davis, MSE = 0.355 and CI = 0.825 improve over DTA-GTOmega (MSE = 0.556, CI = 0.742) by 36.2% and 11.2%. On KIBA, MSE = 0.286 and CI = 0.805 represent a 33.8% MSE reduction and 16.3% CI gain over DTA-GTOmega (MSE = 0.432, CI = 0.692). On BindingDB, MSE = 0.308 and CI = 0.902 correspond to a 79.0% reduction in MSE and a 31.5% improvement in CI over DTA-GTOmega (1.466, 0.686). No test protein overlaps with any training interaction, so these margins directly quantify how well ESM2-derived contact maps transfer to unseen protein families. Contact probabilities encode coevolutionary and structural constraints intrinsic to the sequence. They do not depend on observed interaction labels, and this property plausibly underlies the robust generalisation observed across datasets.

Under the Drug-Target Cold-Start Setting on Davis, our model achieves an MSE of 0.571 and a CI of 0.865, against the best baseline (DTA-GTOmega: MSE = 0.522, CI = 0.628). While MSE is marginally higher than the best baseline, CI improves by 37.7%, indicating substantially stronger affinity-ranking performance. On KIBA, our model achieves MSE = 0.454 and CI = 0.796, outperforming all baselines on both metrics (best baseline: DTA-GTOmega, MSE = 0.619, CI = 0.634), resulting in a 26.7% reduction in MSE and a 25.6% improvement in CI. For BindingDB, the model achieves MSE = 0.549 and CI = 0.812, outperforming all baselines on both metrics, with a 68.7% reduction in MSE over DTA-GTOmega (1.753) and a 26.9% improvement in CI. Taken together, these results indicate that ESM2-derived contact maps, paired with bidirectional cross-attention fusion, generalise more reliably to unseen scaffolds and novel protein families than any evaluated alternative across kinase-focused, heterogeneous, and large-scale benchmarks.

#### 3.2.3 Case studies

Benchmark performance tells only part of the story. To test whether predicted affinities translate to clinically meaningful targets, the model is evaluated on two sets of clinically relevant protein-ligand pairs associated with obesity and cardiovascular disease. Predictions for the case studies are generated using the BindingDB-trained model checkpoint. For each pair, we compare three estimates of binding strength:

i. the model’s predicted affinity, converted to a Gibbs free energy ΔG via

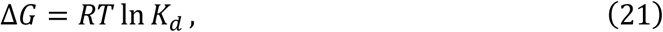

where *R* is the gas constant and *T* is set to 298 K [56] [57].
ii. the AutoDock Vina [58] [59] docking score, obtained under a standard docking protocol, and
iii. experimentally reported ΔG values derived from literature *K*_*i*_ measurements.

### A. Obesity disease

Six protein-ligand pairs spanning four mechanistic classes are evaluated (Table 7): the metabolic phosphatase PTP1B [60], the RNA demethylase FTO [61], the deacetylase SIRT1 [62], the G protein-coupled receptors GLP-1R and MC4R [63][64], and the endocannabinoid receptor CB1 [65]. The model achieves an MAE of 0.93 kcal/mol and a Spearman rank correlation of 0.77 with experimental ΔG values across these six pairs. AutoDock Vina, for which scores were available for five of the six targets, achieves an MAE of 1.84 kcal/mol and a Spearman rank correlation of 0.60, indicating that the sequence-driven model predicts absolute binding strength more accurately and ranks targets more reliably than rigid-receptor docking despite operating without any structural input.

**Table 7.** Comparative Affinity Evaluation of Obesity Across Model Predictions, Docking Scores, and Experimental *ΔG* Values.

| Target | RCSB ID | Drug | $K_d$ predicted<br>(nM) | $\Delta G$ predicted<br>(Kcal/mol) | $\Delta G$ AutoDock<br>(Kcal/mol) | $\Delta G$<br>Experimental<br>(Kcal/mol) |
| --- | --- | --- | --- | --- | --- | --- |
| Protein Tyrosine Phosphatase 1B (PTP1B) | 1C88 | 2-(Oxalyl-Amino)-4,5,6,7-Tetrahydro-Thieno[2,3-C]Pyridine-3-Carboxylic Acid | 1233.10 | -8.06 | -6.30 | -9.05 |
| FTO (Fat mass and obesity-associated protein) | 4IE7 | 4,5-dihydroxy-9,10-dioxo-9,10-dihydroanthracene-2-carboxylic acid | 1472.31 | -7.95 | -9.13 | -7.72 |
| GLP-1 Receptor (GLP1R) | 7LCK | 2-[(4-{6-[(4-cyano-2-fluorophenyl)methoxy]pyridin-2-yl}piperidin-1-yl)methyl]-1-{[(2S)-ssoxetan-2-yl]methyl}-1H-benzimidazole-6-carboxylic acid | 181.55 | -9.19 | -11.72 | -9.68 |
| Melanocortin-4 receptor (MC4R) | 7PIV | Setmelanotide | 9.41 | -10.94 | - | -11.83 |
| CB1 cannabinoid receptor | 6KQI | 2-[(1R,2R,5R)-5-hydroxy-2-(3-hydroxypropyl)cyclohexyl]-5-(2-methyloctan-2-yl)phenol | 21.62 | -10.45 | -10.32 | -12.95 |
| SIRT1 | 5BTR | Resveratrol | 1114.29 | -8.12 | -7.30 | -7.67 |

Five of the six coevolutionary proteins in 1.0 kcal/mol of the experimental value, including FTO (0.23 kcal/mol), Resveratrol/SIRT1 (0.45 kcal/mol), GLP-1R (0 49 kcal/mol), MC4R (0.89 kcal/mol), and PTP1B (0.99 kcal/mol). The close prediction for GLP-1R is particularly notable, despite being a class B GPCR, a receptor family associated with significant conformational flexibility [63], the model achieves a deviation of only 0.49 kcal/mol. GLP-1R possesses a well-structured extracellular ligand-binding domain with a relatively conserved inter-residue contact pattern, which likely enables the ESM2-derived contact map to capture sufficient binding-relevant topology without explicit pocket information. The sole outlier in this case study is CB1 (2.50 kcal/mol), where the model systematically underpredicts binding strength. CB1 is a class A G protein-coupled receptor (GPCR) with an orthosteric binding site located within the transmembrane helical bundle. This lipid-facing cavity exhibits geometry and energetics that are highly dependent on membrane-induced conformational states [65]. Contact maps cannot encode the geometry of transmembrane cavities or lipid context. The underprediction is expected and mechanistically interpretable. Notably, AutoDock Vina, which has access to the receptor structure and binding box, achieves a comparable deviation of 1.63 kcal/mol for CB1, suggesting that the difficulty of this target is intrinsic to the binding mode rather than specific to the sequence-based approach [66].

### B. Cardiovascular diseases

Eleven protein-ligand pairs spanning five therapeutic classes are evaluated (Table 8). ACE [67] inhibitors (Lisinopril, Enalapril, Captopril), HMG-CoA reductase inhibitors [68] (Atorvastatin, Simvastatin), a calcium channel blocker (Diltiazem), an antiarrhythmic (Flecainide), an anticoagulant (Rivaroxaban), a cardiac glycoside binding [69] (Digoxin), an endothelin receptor antagonist [70] (Bosentan), and an aldosterone receptor antagonist (Spironolactone). The model achieves an MAE of 1.76 kcal/mol across these eleven pairs, compared to 2.81 kcal/mol for AutoDock Vina. The cardiovascular set presents a more challenging evaluation than the obesity set, primarily because it contains a higher proportion of targets whose dominant binding contributions involve interaction features absent from the model’s input representation.

**Table 8.** Comparative Affinity Evaluation of Cardiovascular diseases Across Model Predictions, Docking Scores, and Experimental ΔG Values.

| Target | RCSB ID | Drug | $K_d$ predicted<br>(nM) | $\Delta G$ predicted<br>(Kcal/mol) | $\Delta G$ AutoDock<br>(Kcal/mol) | $\Delta G$ Experimental<br>(Kcal/mol) |
| --- | --- | --- | --- | --- | --- | --- |
| Angiotensin-Converting Enzyme (ACE) | 1O86 | Lisinopril | 210.37 | -9.10 | -7.53 | -11.34 |
| Angiotensin-Converting Enzyme (ACE) | 1UZE | Enalapril | 326.58 | -8.84 | -7.57 | -12.07 |
| Angiotensin-Converting Enzyme (ACE) | 4C2P | Captopril | 134.58 | -9.37 | -5.95 | -11.96 |
| HMG-CoA Reductase (HMGCR) | 1HWK | Atorvastatin | 267.30 | -8.96 | -8.98 | -11.19 |
| HMG-CoA Reductase (HMGCR) | 1HW9 | Simvastatin | 22.59 | -10.42 | -7.53 | -10.56 |
| L-type Calcium Channel Alpha-1C (CACNA1C) | 6JPB | Diltiazem | 93.75 | -9.58 | -6.25 | -8.75 |
| Voltage-gated Sodium Channel Nav1.5 (SCN5A) | 6UZ0 | Flecainide | 306.90 | -8.88 | -8.15 | -7.07 |
| Coagulation Factor Xa (F10) | 5VOF | Rivaroxaban | 132.13 | -9.38 | -7.30 | -11.55 |
| Na<sup>+</sup>/K<sup>+</sup>-ATPase Alpha-1 (ATP1A1) | 7DDH | Digoxin | 85.70 | -9.64 | -11.75 | -10.05 |
| Endothelin Receptor Type B | 5XPR | Bosentan | 827.94 | -8.29 | -9.02 | -8.82 |
| Aldosterone Receptor (NR3C2) | 2AB2 | Spironolactone | 988.55 | -8.19 | -9.68 | -11.32 |

Four of the eleven pairs fall within 1.0 kcal/mol of the experimental value: Simvastatin (0.14 kcal/mol), Digoxin (0.41 kcal mol), Bosentan (0.53 kcal/mol), and Diltiazem (0.83 kcal/mol). The closest prediction for Digoxin is worth singling out.

Na⁺K⁺- ATPase [69] is a large multi-pass transmembrane protein, exactly the kind of target where sequence-only methods are expected to struggle, yet the predicted affinity lands within 0.41 kcal/mol of the experimental value. That the model recovers this accurately without any structural input suggests that ESM2-derived contact maps capture binding-relevant spatial organisation even for membrane proteins, provided the drug-binding domain is sufficiently conserved at the sequence level.

The remaining six pairs, all with deviations exceeding 2 0 kcal/mol, cluster into two mechanistically distinct classes. The first is zinc-coordinating metalloenzymes, all three ACE inhibitors (2.24-3.23 kcal/mol) and Rivaroxaban (2.17 kcal/mol) show systematic underprediction of binding strength. Zinc coordination is not a contact it is a directional, high-energy interaction between a chelating functional group and a specific metal ion [67] whose position and geometry have to be known precisely to model correctly. Neither residue-level descriptors nor ESM2 contact probabilities carry that information, and there is no way to recover it from the sequence alone. The model has no knowledge of where on the protein the ligand actually binds. For ACE inhibitors, which share a well-defined catalytic site, the blind spot compounds the model cannot weight residues by their proximity to the active centre, and the systematic underestimation seen across the entire inhibitor class follows directly from that absence. AutoDock Vina, despite having access to the receptor structure [66] and binding pocket, performs substantially worse on this class (errors of 3.97-6.01 kcal/mol), consistent with the known difficulty of scoring zinc metalloprotease interactions with empirical docking functions. The second class covers receptors whose binding cavities undergo substantial rearrangement upon ligand entry. Spironolactone binding to the mineralocorticoid receptor is a case in point, the receptor remodels around the ligand rather than accepting it into a preformed pocket, and a static contact map has no way to represent that process. The predicted affinity of 3.13 kcal/mol reflects that the model is working from a frozen snapshot of a protein that does not stay still.

For within-class potency ranking, the operationally relevant capability for lead optimisation, the model correctly identifies Simvastatin as a strong HMGCR binder and Captopril as the most potent ACE inhibitor in the predicted ordering. The model misranks Enalapril [71] relative to Lisinopril within the ACE class. This interpretation warrants verification against the original experimental source for this *K*_*i*_ value.

Across all 17 pairs, the model achieves a combined MAE of 1.46 kcal/mol and a Spearman rank correlation of 0.50 with experimental ΔG values, compared to an MAE of 2.51 kcal/mol and a Spearman rank correlation of 0.01 for AutoDock Vina. The contrast between the two case studies is itself informative. The obesity set (MAE 0.93, Spearman 0.77) is handled substantially better than the cardiovascular set (MAE 1.76), and the difference is explained almost entirely by the higher concentration of zinc-coordinating targets in the cardiovascular panel. This pattern draws a reasonably clear boundary around what XAttn-DTA can and cannot do. Where binding affinity is driven by structural and physicochemical complementarity that leaves a detectable signature at the sequence level, predictions are reliable. Where it is dominated by zinc coordination [67] or membrane-context-dependent cavity geometry [65], they are not, and no purely sequence-driven approach without explicit binding site priors will do much better. Incorporating metal-binding residue annotations or binding-pocket embeddings as additional input features is the most direct path to addressing these limitations in future work.

## 4. Future Work

The benchmark evaluations and case studies revealed clear limitations that point to specific areas for future research. The Davis drug-target cold-start results show that absolute regression accuracy degrades when neither modality is observed during training. The current contact map representations may lack sufficient structural priors for completely unseen protein families, motivating the incorporation of confidence-weighted or distance-aware contact representations from ESM2 that more explicitly encode positional uncertainty. The protein graph has no dynamics. Every contact map in the pipeline is a single frozen snapshot, one conformation, one threshold, and anything the protein does beyond that is simply not represented. Conformational flexibility, allosteric shifts, and residue mobility near the binding site, none of it survives the thresholding step that produces the graph. Representing proteins as ensembles of predicted contact maps or as dynamic graph structures that capture residue mobility could better address induced-fit binding scenarios beyond the reach of a fixed-topology graph. Atom-level message passing has a range limit. The GNN sees bonds and immediate neighbourhoods well enough. Still, scaffold-level generalisation, the kind of cold-start evaluation specifically probed, requires chemical reasoning at a coarser granularity than individual atoms provide. Fragment vocabularies and pretrained molecular transformers both work at that granularity. Either could sit upstream of the GNN. The existing architecture would not need to change anywhere else. The single-task objective is the most consequential limitation in practice. Affinity alone is not what anyone actually optimizes for in a drug discovery campaign toxicity, selectivity, and half-life matter just as much and often more. A shared encoder trained jointly across these endpoints would receive more gradient signal per molecule and might generalise better, particularly where labelled affinity data is scarcest. We have not tested this, and it is not a trivial extension, but it is the direction we would pursue first.

## 5. Conclusion

XAttn-DTA was developed based on the hypothesis that integrating graph-based drug encoding, ESM2-derived protein contact maps, and bidirectional cross-attention fusion produces a more generalizable affinity predictor than existing methods. Under warm-start and cold-start settings, XAttn-DTA demonstrates consistent improvements in mean squared error (MSE) and concordance index (CI) across respective datasets, with one notable exception, absolute regression accuracy under the Davis drug-target cold-start condition. In this scenario, a maximal distribution shift reveals a tension between ranking reliability and egression precision that ranking metrics alone do not capture. Evaluation on BindingDB, which is larger and more chemically diverse than either Davis or KIBA, indicates that the observed generalization gains are not merely a consequence of the kinase-heavy training distribution, these gains persist at scale. Qualitative case studies on obesity and cardiovascular targets show that predicted affinities generally match experimental results across different types of therapies. It is important because these targets differ significantly from one another. Collectively, these findings show that ESM2-derived structural representations, when fused via cross-modal attention, provide the model with a substantive advantage when encountering novel scaffolds or unfamiliar protein families. Since therapeutic targets with limited structural coverage are common and structure-based methods are impractical at the scale required for early drug discovery, a model that generalizes effectively without relying on resolved structures addresses a significant unmet need.

## Acknowledgements

The authors would like to express their sincere gratitude to all individuals who contributed their time, expertise, and support throughout this research.

